# Red Sea *Synechococcus* exhibits greater resilience to UV-B radiation than pico-eukaryotic phytoplankton

**DOI:** 10.1101/2025.02.18.638780

**Authors:** Sebastian Overmans, Susana Agustí

## Abstract

UV-B radiation can be a significant stressor to primary producers in tropical seas, particularly to organisms residing close to the surface. We investigated the UV-B sensitivity of natural pico-phytoplankton communities in the Red Sea, one of the warmest seas on Earth. The communities were incubated either in full sunlight or under UV-B-reduction conditions. The mean lethal UV-B radiation dose (LRD_50_) of natural *Synechococcus* was 57 kJ to 73% non-detectable compared to 8 KJ to 45 % non-detectable for pico-eukaryotes. UV-B-induced *F*_v_/*F*_m_ change (Δ*F*_v_/*F*_m_) in natural pico-phytoplankton communities ranged from -0.22 to 0.06 (mean: -0.08). Changes in *Synechococcus* growth under UV-B-reduction were more pronounced in cooler waters. The dominant pico-phytoplankton in the Red Sea, *Synechococcus*, exhibited a strong resistance to UV-B, highlighting their exceptional adaptation to the local conditions. Our results underscore the importance of considering the UV-B sensitivity of natural phytoplankton communities in the context of global change, which may differ from those of cultured strains.

## 2. Introduction

Marine photoautotrophs are photosynthetic organisms that are restricted to the sunlit zone of the ocean, where they rely on photosynthetically active radiation (PAR: 400–700 nm) to meet their metabolic needs. However, solar ultraviolet radiation (UVR: 200–400 nm) can have predominantly adverse effects on marine biota, particularly on primary producers such as phytoplankton. In the marine realm, there are two UVR waveband spectra, UV-A (320–400 nm) and UV-B (280–320 nm), with the latter considered more harmful to phytoplankton due to its shorter, more energetic wavelengths (Häder, 1997; Vernet, 2000). Although UV-B is generally attenuated in the water column, it can reach considerable depths in shallow or oligotrophic waters where chromophoric dissolved organic matter (CDOM), the primary UVR attenuator, is minimal. This can represent a crucial abiotic stressor for marine biota. For example, exposure of phytoplankton to UV-B radiation can lead to severe effects such as elevated levels of oxidative stress (Zhang et al., 2017), decreased photosynthetic performance (Cai et al., 2017; Conan et al., 2008; Garcia-Corral et al., 2017; Li et al., 2015; Shi et al., 2017), impaired growth (Nahon et al., 2010), increased DNA damage (Häder and Gao, 2015) and higher mortality rates (Llabres and Agusti, 2006; Llabres et al., 2010). It is well known the detrimental effects of UV-B exposure on cells are diminished by exposure to moderate intensities of UV-A radiation (Häder et al., 2015). Specifically, UV-A wavelengths aid in the photoreactivation of DNA and initiates cellular repair processes (Li et al., 2011; Quesada et al., 1995; Sinha and Häder, 2014), which highlights the importance of using ecologically relevant spectral compositions during experiments that aim to evaluate UV-B impacts on phytoplankton.

Phytoplankton cells have evolved protective strategies to mitigate the damage caused by UV-B radiation. For example, many species, especially dinoflagellates, produce photoprotective compounds known as mycosporine-like amino acids (MAAs) that can absorb or scatter UV-B wavelengths and prevent them from reaching sensitive organelles (Joshi et al., 2018; Laurion et al., 2004). MAAs can also function as antioxidants(Browne et al., 2023; Gauthier et al., 2020), which can counteract the effects of reactive oxygen species (ROS) that are produced by UV-B exposure (Costa et al., 2021; Huang et al., 2018; Suh et al., 2003). Another example is the pigment scytonemin, which is synthesized by some cyanobacteria and can help protect against UV damage (Rastogi et al., 2013). However, the effectiveness of these UV-absorbing compounds is highly dependent on factors such as the density of pigments or cell size, which affect the path length that UV-B wavelengths must traverse (Garcia-Pichel, 1994; Laurion et al., 2004).

Cell size has been suggested to be a key factor in determining the sensitivity of phytoplankton to UV-B radiation (Boelen et al., 2000; Hader et al., 2007). As cell size decreases, it becomes increasingly difficult for the cells to accumulate a sufficiently thick layer of photoprotective compounds, making them more susceptible to damage (Garcia-Pichel, 1994). The role of cell size in determining the susceptibility of phytoplankton to UV-B damage has been demonstrated for cyanobacteria such as *Synechococcus* and *Prochlorococcus*, where smaller *Prochlorococcus* cells were more severely affected by UV radiation, leading to higher mortality rates (Llabres and Agusti, 2006; Llabres et al., 2010). Similarly, a comparison of 12 Antarctic diatoms found that smaller diatoms (e.g., *Chaetoceros* sp.), with larger surface area-to-volume ratios, sustained more DNA damage and had lower survival rates than larger diatoms (e.g., *Coscinodiscus* sp.) under UV exposure (Karentz et al., 1991). This trend was also observed in high light conditions, where smaller diatoms were more strongly inhibited by UV radiation than larger ones (Wu et al., 2015).

Several studies have investigated the factors that determine the sensitivity of phytoplankton to UV radiation. Some studies have suggested that UV sensitivity is related to the bio-optical properties of phytoplankton rather than their cell size or chlorophyll concentration (Figueroa et al., 1997). As bio-optical and cellular properties, such as cell morphology and the position of organelles, vary considerably between phytoplankton taxa, UV effects are likely taxon-dependent (Jin et al., 2022; Laurion and Vincent, 1998). For example, diatoms, which have siliceous cell walls, may be vulnerable to UVR because their organelles are essentially unprotected against UV exposure (Karentz et al., 1991). Other phytoplankton taxa, such as pico-cyanobacteria, lack the ability to utilize mycosporine-like amino acids (MAAs) as optical barriers against UVR due to their small size. Instead, they have developed various defence strategies, such as multiple copies of their genome, adaptive mutagenesis, ROS-quenching enzymes, and non-enzymatic antioxidant compounds, to minimize UV-induced damage (Kumar et al., 2016; Prabha and Kulandaivelu, 2002).

In the Red Sea, the phytoplankton community is dominated by pico-cyanobacteria, especially *Synechococcus* and *Prochlorococcus* (Kheireddine et al., 2017; Stambler, 2006), due to the oligotrophic state of the basin, which favours small-sized autotrophs (Johnson et al., 2006; Partensky et al., 1999). Phytoplankton biomass in the Red Sea is generally low, with chlorophyll concentrations < 0.8 mg Chl m^−3^ (López-Sandoval et al., 2019), except in the south where nutrients are continually replenished by the influx of waters from the Gulf of Aden (Raitsos et al., 2013). Consequently, chlorophyll concentrations can reach 4 mg Chl m^−3^ in the southern Red Sea during spring bloom events.

The Red Sea displays a prominent north-south gradient in sea surface temperature (SSTs) and experiences extreme summertime SST maxima, with values up to 34°C in open waters and 35°C in shallow coastal ecosystems (Chaidez et al., 2017; Garcias-Bonet and Duarte, 2017; Genevier et al., 2019). These features are particularly relevant in the context of UV sensitivity, as temperature has been shown to play a crucial role in governing the vulnerability of phytoplankton to UV radiation. Temperature affects the rate and efficiency of enzymatic conversions and cellular repair mechanisms (Bouchard et al., 2006; Halac et al., 2014; Helbling et al., 2011), while extreme temperatures can intensify the damaging effects of high irradiance or vice versa (Giordanino et al., 2011). Therefore, the pronounced temperature gradient and extreme summer temperatures in the Red Sea may significantly impact the UV sensitivity of phytoplankton communities in this region. In addition, phytoplankton blooms in the Red Sea during late spring and early summer coincide with incident UV irradiance peaks (Overmans and Agusti, 2020) and the water column reaches its highest transparency to ultraviolet radiation (Dishon et al., 2012b; Overmans and Agusti, 2019, 2020). Consequently, phytoplankton assemblages experience extreme UV-B exposure even at considerable depths within the upper layer of the euphotic zone.

In this study, our aim was to evaluate the sensitivity to UV-B radiation of natural phytoplankton communities along a latitudinal gradient in the Red Sea. Specifically, we assessed the extent to which UV-B exposure modulates the growth or decay of the phytoplanktonic organisms and evaluated the impacts of UV-B exposure on the photosystem II by measuring the extent of photoinhibition based on maximum photochemical efficiency (*F*_v_/*F*_m_) under different UV-B exposures. Our results provide novel insights into the differential tolerance of various Red Sea phytoplankton taxa to solar UV-B, which can be used to compare their UV-B sensitivities against those of other marine phytoplankton around the globe.

## 3. Materials & Methods

### 3.1 Natural phytoplankton community experiments

Between October 2016 and March 2018, a total of 11 experiments were conducted to evaluate the sensitivity of phytoplankton communities along the latitudinal axis of the Red Sea to UV-B radiation (Fig. 1 and Table 1). All field experiments were approved by KAUST, and the necessary sampling permits were obtained from the Saudi General Directorate of the Border Guard through the Coastal and Marine Resources Core Lab (CMR). At each station, seawater was collected from a depth of 5 m using a Niskin bottle in the morning (8–9 am), pre-filtered (100 µm pore size), and then transferred into 12 bottles (four borosilicate glass, four quartz glass, four black) with a volume of 300 ml each. The full sunlight (FS) treatment was applied to the transparent quartz glass bottles that transmit PAR + UV-A + UV-B radiation, while the UV-B-reduction (-UV-B) treatment was applied to the borosilicate glass bottles that transmit almost exclusively PAR + UV-A (See Suppl. Fig. 1). The bottles were then placed upside down into a shallow (∼ 50 cm depth) open-top incubator supplied with flow-through surface seawater to maintain in situ temperatures. Incubations were carried out under natural solar radiation, but a shade cloth (∼76% light transmission, Suppl. Fig. 1) was installed around the incubator to create UV-B conditions more similar to those in the first few meters of the water column. All samples were incubated for 9 hours (9 am–6 pm), and a 15 ml sample was collected from each bottle at 9 am (T_0_), 12 pm (T_1_), and 6 pm (T_2_) to measure the abundance of cyanobacteria and pico-eukaryotes, and to evaluate the photosynthetic efficiency (*F*_v_/*F*_m_) of the phytoplankton community. The range of UV-B exposure during the incubation periods was between 22.70 and 30.72 kJ m^−2^ (Table 1). The incident PAR and UV-B irradiances were recorded underneath the shade cloth using a PMA2100 data-logging radiometer (Solar Light, USA) fitted with a PAR (400–700 nm, model PMA2132-WP) and a UV-B (280–320 nm, model PMA2106-WP) sensor.

**Figure 1.**
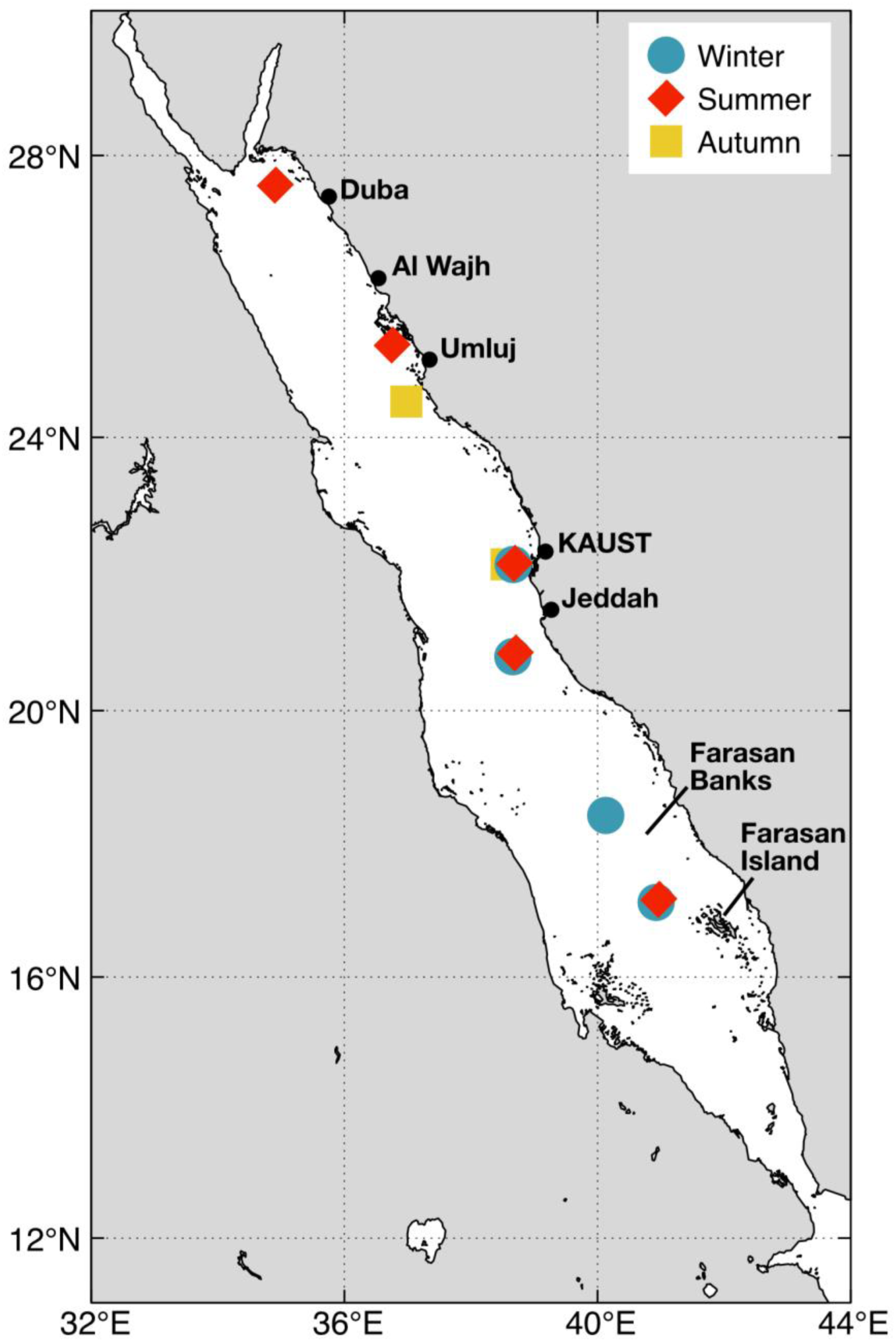
Location of the sampling stations in the Red Sea where phytoplankton UV experiments were performed as part of three research cruises: *Winter CCF* (blue circles), *Summer CCF* (red diamonds), and *Threats cruise* (yellow squares).

**Table 1.**
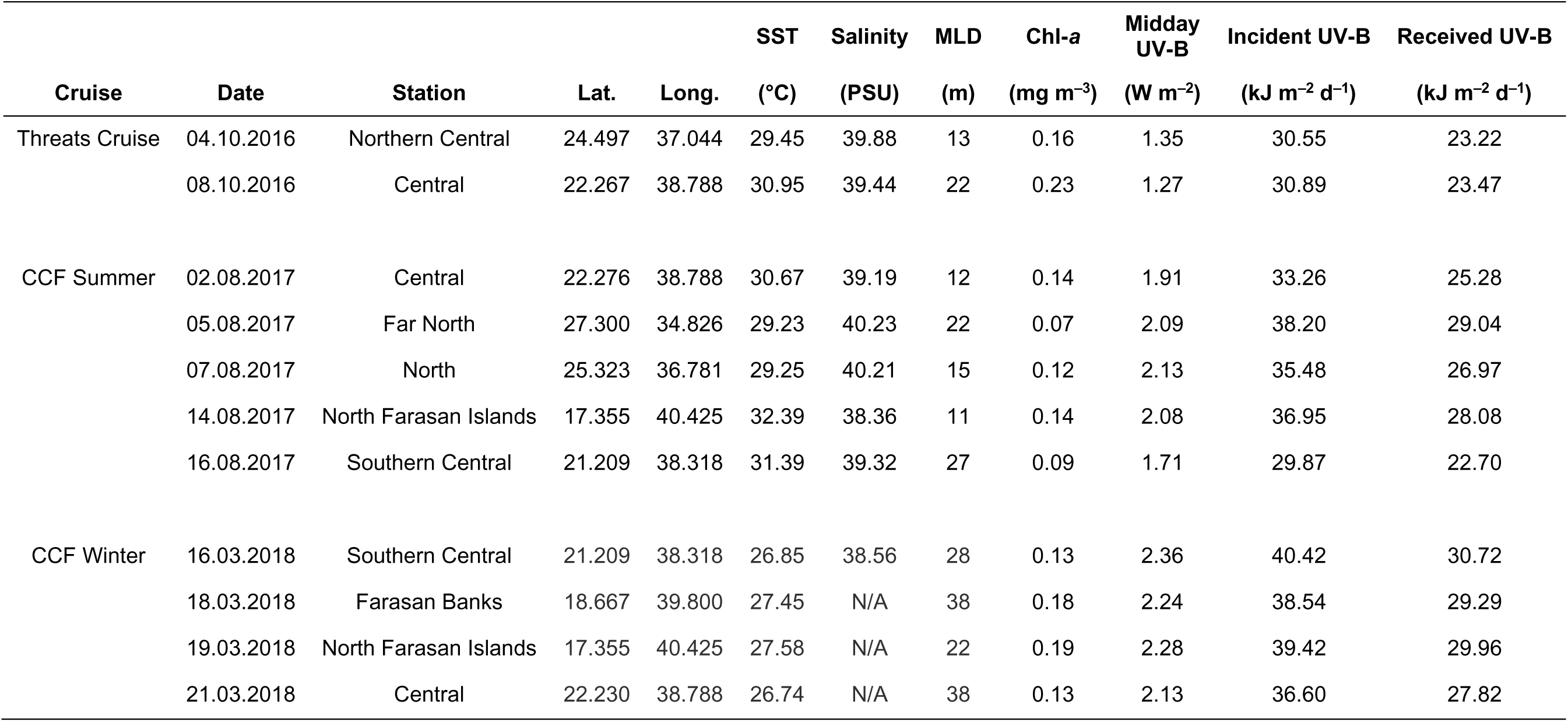
Location and environmental conditions of the Red Sea sampling stations during phytoplankton incubation experiments performed as part of three research cruises between October 2016 and March 2018. Received UV-B exposures refer to the exposures measured inside the incubator, while incident exposures above the shade cloth were calculated by using a factor of 0.76, representing a 76% light transmission of the shade cloth. Mixed layer depth (MLD) has been adapted from Overmans and Agusti (2019).

|  |  |  |  |  | SST | Salinity | MLD | Chl-a | Midday<br>UV-B | Incident UV-B | Received UV-B |
| --- | --- | --- | --- | --- | --- | --- | --- | --- | --- | --- | --- |
| Cruise | Date | Station | Lat. | Long. | (°C) | (PSU) | (m) | (mg m<sup>-3</sup>) | (W m<sup>-2</sup>) | (kJ m<sup>-2</sup> d<sup>-1</sup>) | (kJ m<sup>-2</sup> d<sup>-1</sup>) |
| Threats Cruise | 04.10.2016 | Northern Central | 24.497 | 37.044 | 29.45 | 39.88 | 13 | 0.16 | 1.35 | 30.55 | 23.22 |
|  | 08.10.2016 | Central | 22.267 | 38.788 | 30.95 | 39.44 | 22 | 0.23 | 1.27 | 30.89 | 23.47 |
| CCF Summer | 02.08.2017 | Central | 22.276 | 38.788 | 30.67 | 39.19 | 12 | 0.14 | 1.91 | 33.26 | 25.28 |
|  | 05.08.2017 | Far North | 27.300 | 34.826 | 29.23 | 40.23 | 22 | 0.07 | 2.09 | 38.20 | 29.04 |
|  | 07.08.2017 | North | 25.323 | 36.781 | 29.25 | 40.21 | 15 | 0.12 | 2.13 | 35.48 | 26.97 |
|  | 14.08.2017 | North Farasan Islands | 17.355 | 40.425 | 32.39 | 38.36 | 11 | 0.14 | 2.08 | 36.95 | 28.08 |
|  | 16.08.2017 | Southern Central | 21.209 | 38.318 | 31.39 | 39.32 | 27 | 0.09 | 1.71 | 29.87 | 22.70 |
| CCF Winter | 16.03.2018 | Southern Central | 21.209 | 38.318 | 26.85 | 38.56 | 28 | 0.13 | 2.36 | 40.42 | 30.72 |
|  | 18.03.2018 | Farasan Banks | 18.667 | 39.800 | 27.45 | N/A | 38 | 0.18 | 2.24 | 38.54 | 29.29 |
|  | 19.03.2018 | North Farasan Islands | 17.355 | 40.425 | 27.58 | N/A | 22 | 0.19 | 2.28 | 39.42 | 29.96 |
|  | 21.03.2018 | Central | 22.230 | 38.788 | 26.74 | N/A | 38 | 0.13 | 2.13 | 36.60 | 27.82 |

### 3.2 Flow cytometry

Cell abundances in natural phytoplankton communities were determined flow-cytometrically at the onset of experiments and twice daily thereafter. For the phytoplankton community experiments, a CyFlow Space flow cytometer (Sysmex Corporation, Japan) equipped with a 96-well plate autoloading station was used. Each sample was pipetted into a separate well of a 96-well microplate (500 µl in duplicates) and loaded into the autosampler. Sample acquisitions automatically stopped once the instrument sampled exactly 150 µl from each well. Two technical replicates per biological replicate (i.e., incubation bottle) were analyzed. The number of recorded phytoplankton cells was used to calculate cell abundance (in cells ml^−1^). FCS Express 6 Flow (De Novo Software, USA) was used for all post-acquisition analyses and population clustering.

### 3.3 Growth rate (µ), half-life time (t_1/2_), and LRD_50_ calculations

The growth rate (µ, in h^−1^) of each natural community sample was determined as the slope of the linear regression between the natural logarithm (ln) of the change in cell abundance with time (in h). Absolute changes in growth rates due to UV-B exposure (Δµ) were calculated as the difference between µ of the FS and -UV-B treatments (Eqn 1):

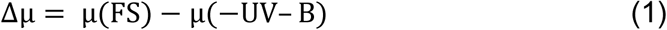

The half-life times (t_1/2_, in h) of populations, i.e., the time required to reduce the initial cell density by 50%, were calculated using the formula described by Llabres and Agusti (2006) (Eqn 2):

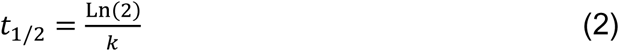

where the decay rate *k* (h^−1^) is calculated as the linear regression slope between the natural logarithm of the cell abundance and time. Similarly, the 50% lethal radiation dose (LRD_50_, in kJ m^−2^), defined as the UV-B exposure required to decrease the initial cell density by half, was calculated using k, which refers to the population’s decay rate (in per kJ m^−2^), determined as the slope of the relationship between the natural logarithm of the cell abundance and UV-B radiation exposure (in kJ m^−2^). For each phytoplankton group, the theoretical daily population decay (in % of initial concentration) during the months of July and January in 0.5, 2 and 8 m depth (*z*) was calculated based on the decay rate *k* (in per kJ m^−2^) and using the daily UV-B exposures in those depths (Iz, in kJ m^−2^ d^−1^) reported by Overmans and Agusti (2020):

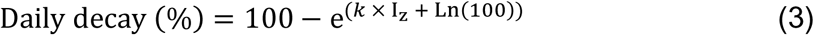

### 3.4 Maximum photochemical efficiency (*F*_v_/*F*_m_)

The inhibition of the photosynthetic apparatus of natural communities was evaluated by measuring variable chlorophyll fluorescence of photosystem II (PSII) using a Pulse Amplitude Modulation (PAM) fluorometer (Phyto-PAM; Heinz Walz GmbH, Germany) equipped with an emitter-detector-unit (Phyto-ED; Heinz Walz GmbH, Germany). Before each measurement, the samples were dark-adapted for 15 min at the respective experimental in situ incubation temperatures (27–32 °C) to ensure all PSII reaction centres were open. For each sample, the signal amplitude was adjusted, and a filtered (0.2 μm) blank sample was used to set an automatic baseline adjustment before two single-turnover measurements were recorded per sample flask. The maximum photochemical efficiency (*F*_v_/*F*_m_) was determined using the following equation (Eqn 4):

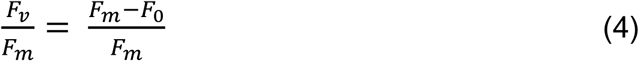

where *F*_m_ and *F*_0_ are the maximal and minimal PSII fluorescence of dark-acclimated phytoplankton cells, respectively (Baker, 2008). Absolute changes in *F*_v_/*F*_m_ due to UV-B exposure (Δ *F*_v_/*F*_m_) were calculated as the difference between *F*_v_/*F*_m_ of the full sunlight and -UV-B treatments. All data were recorded and analyzed using the software PhytoWin (v3.t54).

### 3.5 Statistical analysis

Kruskal-Wallis tests were performed followed by Wilcoxon each pair post hoc tests to identify significant variations (*p* < 0.05) of cell abundances during the cruise experiments between the three radiation treatments. The mean growth rates and *F*_v_/*F*_m_ ratios of the FS and -UV-B treatment were determined at each sampling point during the cruise experiments using individual Student’s *t*-tests. Pearson’s correlation coefficients (*r*) were calculated to determine the strength of association of various environmental parameters with Δμ of picophytoplankton and the community Δμ *Fv/Fm*. Data analysis and visualization were carried out using either JMP Pro 14.1.0 (SAS Institute Inc., Cary, North Carolina, USA) or GraphPad Prism 8.3.0 (GraphPad Software Inc., CA, USA).

## 4. Results

### 4.1 Population dynamics and UV-B-induced cell decay

Considerable variation in initial cell concentration and UV-B-induced mortality rates were observed between different taxa and natural phytoplankton communities sampled at different locations and seasons. At T0, *Synechococcus* cell concentrations ranged from 919–26,493 cells ml^−1^ (Fig. 2) and pico-eukaryotes concentrations ranged from 42– 1,465 ml^−1^ (Fig. 3). Both taxa showed the lowest cell concentrations during summer and the highest in autumn. Mean cell concentrations of *Synechococcus* in the dark bottles were generally higher at T2 compared to T0 (Fig. 2), while for pico-eukaryotes, this trend was observed in only four out of 11 experiments (Fig. 3).

**Figure 2.**
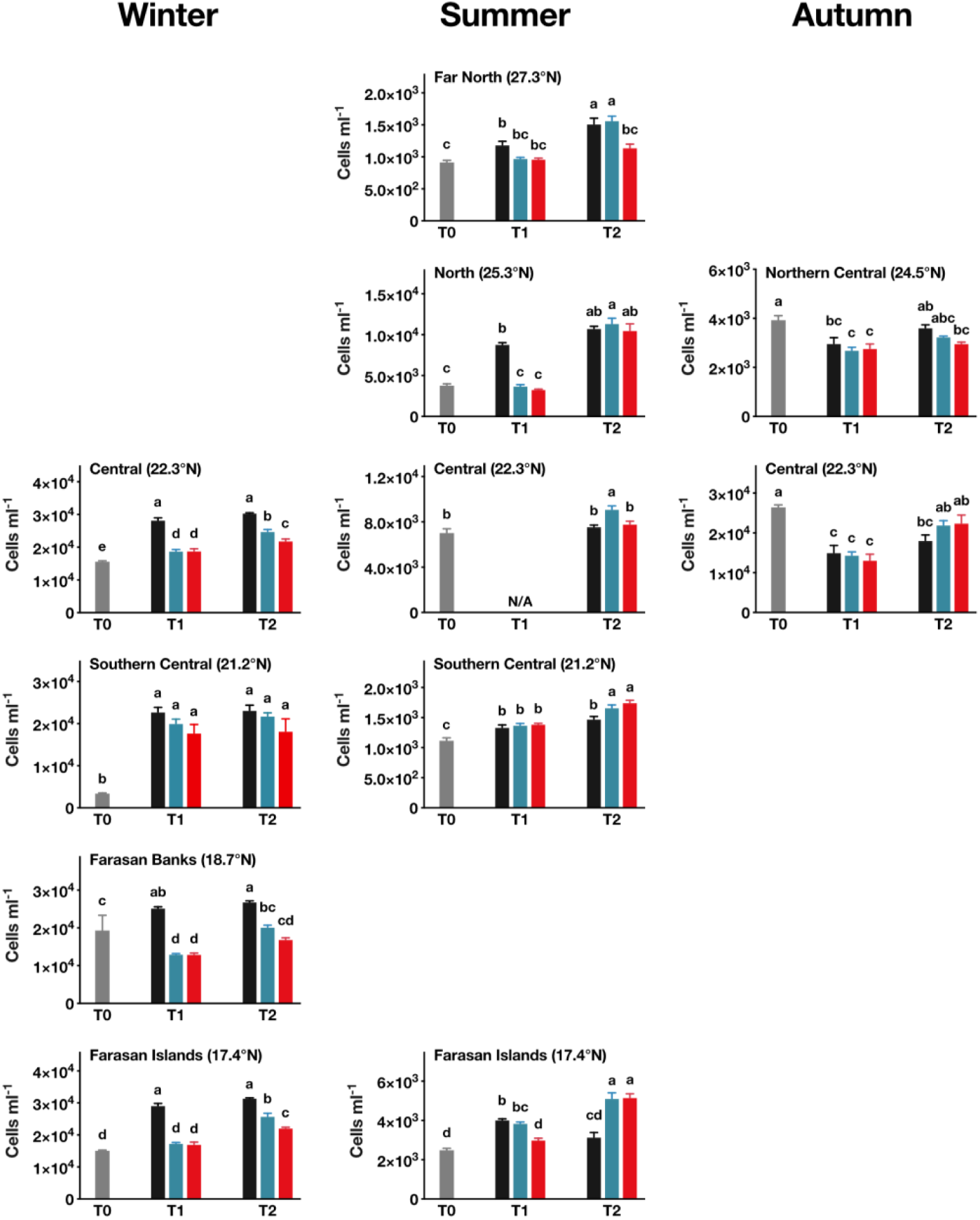
*Synechococcus* sp. cell abundance (cells ml^−1^) during the research cruise experiments in winter (left column), summer (middle column) and autumn (right column) from the northernmost station to the Farasan Islands in the far south of the basin (bottom row). T0 measurements are shown in grey while the colors of the other bars indicate the different treatments: complete darkness (in black), UVB-reduction (-UVB, in blue), full sunlight (FS, in red). Bars that do not share the same letter represent significant differences at a level of *p* < 0.05. In cases where no letters are displayed, none of the results were significantly different from each other.

**Figure 3.**
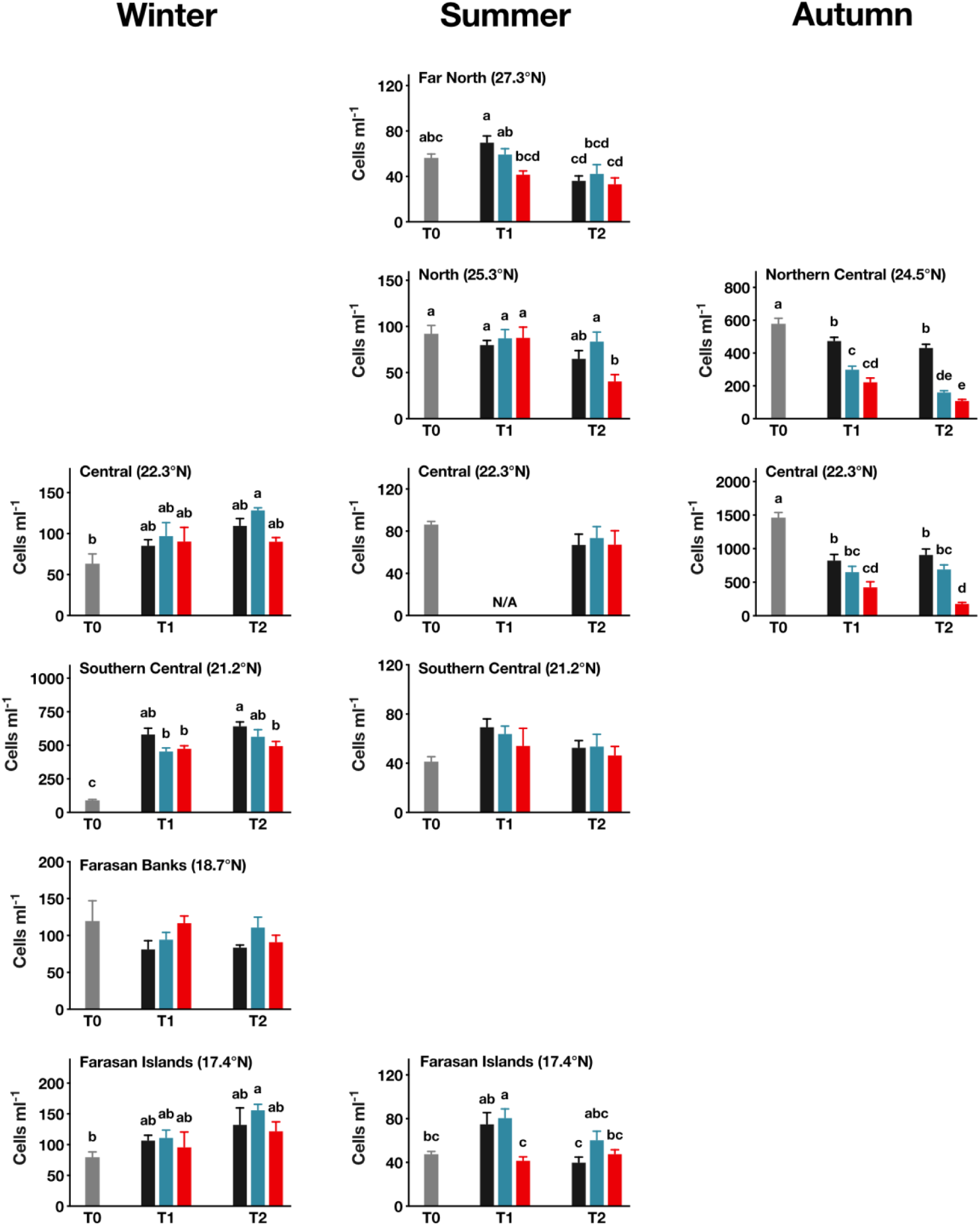
Pico-eukaryote cell abundance (cells ml^−1^) during the research cruise experiments in winter (left column), summer (middle column) and autumn (right column) from the northernmost station to the Farasan Islands in the far south of the basin (bottom row). For a description of the color-coding, see Fig. 2.

Growth rates of *Synechococcus* ranged from -0.031 to 0.186 h^−1^ (mean: 0.044 h^−1^) under full sunlight, and from -0.022 to 0.206 h^−1^ (mean: 0.056 h^−1^) in the UV-B-reduction treatment (Table 2). Pico-eukaryotes had lower growth rates than *Synechococcus*, ranging from -0.231 to 0.188 h^−1^ (mean: -0.030 h^−1^) under full sunlight, and from -0.141 to 0.203 h^−1^ (mean: 0.011 h^−1^) in the -UV-B treatment (Table 2). However, the reduction of UV-B did not significantly increase mean growth rate for either phytoplankton group. *Synechococcus* exhibited significantly decreased cell concentrations under full sunlight compared to the UV-B reduction treatment in winter in the central Red Sea (t(DOF: 21)= –3.98, *p* < 0.001) and Farasan Islands (t(21)= –4.95, *p* < 0.001), as well as in summer in the northern (t(21)= –5.51, *p* < 0.001) and central (t(12)= –3.36, *p* = 0.0057) areas (Fig. 2). Pico-eukaryotes showed a significant UV-B-induced reduction in cell concentration during experiments in the North Red Sea in summer (t(21)= –3.59, *p* = 0.0017) and the central Red Sea in autumn (t(14)= –5.17, *p* < 0.001) (Fig. 3).

**Table 2.**
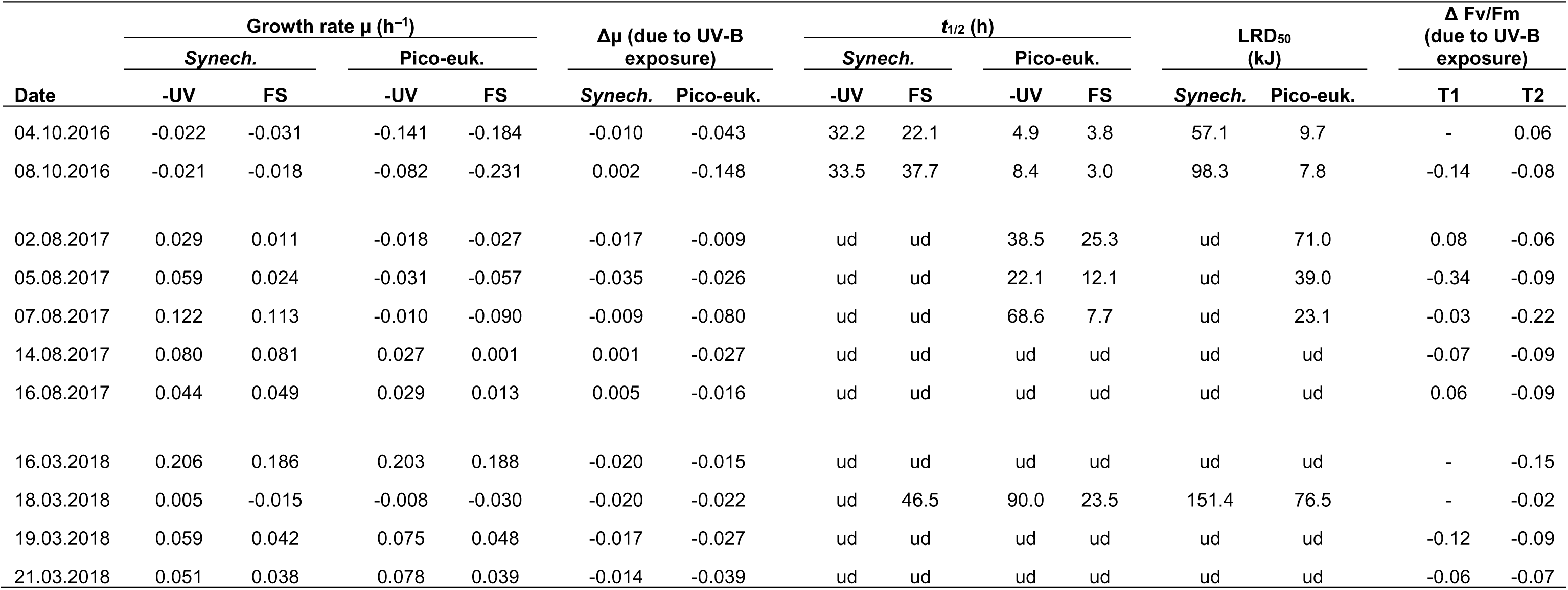
Growth rates (µ) and maximum photochemical efficiency (*F*_v_/*F*_m_) of natural phytoplankton, measured during the cruise experiments. Samples were exposed to either full sunlight (FS) or kept under reduced UV-B conditions (-UV). Also displayed are the calculated half-life times (*t*_1/2_) and lethal radiation doses (LRD_50_, in kJ UV-B). ud = undetectable (i.e., t_1/2_ or LRD_50_ could not be calculated due to a positive growth rate). *F*_v_/*F*_m_ was measured at 12 pm (T1) and 6 pm (T2). Dashes (-) indicate instances where Δ*F*_v_/*F*_m_ could not be determined due to a minimal fluorescence signal.

| Date | Growth rate $\mu$ (h<sup>-1</sup>) | | | | $\Delta\mu$ (due to UV-B exposure) | | $t_{1/2}$ (h) | | | | LRD<sub>50</sub> (kJ) | | $\Delta F_v/F_m$ (due to UV-B exposure) | |
| --- | --- | --- | --- | --- | --- | --- | --- | --- | --- | --- | --- | --- | --- | --- |
|  | *Synech.* |  | Pico-euk. |  |  |  | *Synech.* |  | Pico-euk. |  |  |  |  |  |
|  | -UV | FS | -UV | FS | *Synech.* | Pico-euk. | -UV | FS | -UV | FS | *Synech.* | Pico-euk. | T1 | T2 |
| 04.10.2016 | -0.022 | -0.031 | -0.141 | -0.184 | -0.010 | -0.043 | 32.2 | 22.1 | 4.9 | 3.8 | 57.1 | 9.7 | - | 0.06 |
| 08.10.2016 | -0.021 | -0.018 | -0.082 | -0.231 | 0.002 | -0.148 | 33.5 | 37.7 | 8.4 | 3.0 | 98.3 | 7.8 | -0.14 | -0.08 |
| 02.08.2017 | 0.029 | 0.011 | -0.018 | -0.027 | -0.017 | -0.009 | ud | ud | 38.5 | 25.3 | ud | 71.0 | 0.08 | -0.06 |
| 05.08.2017 | 0.059 | 0.024 | -0.031 | -0.057 | -0.035 | -0.026 | ud | ud | 22.1 | 12.1 | ud | 39.0 | -0.34 | -0.09 |
| 07.08.2017 | 0.122 | 0.113 | -0.010 | -0.090 | -0.009 | -0.080 | ud | ud | 68.6 | 7.7 | ud | 23.1 | -0.03 | -0.22 |
| 14.08.2017 | 0.080 | 0.081 | 0.027 | 0.001 | 0.001 | -0.027 | ud | ud | ud | ud | ud | ud | -0.07 | -0.09 |
| 16.08.2017 | 0.044 | 0.049 | 0.029 | 0.013 | 0.005 | -0.016 | ud | ud | ud | ud | ud | ud | 0.06 | -0.09 |
| 16.03.2018 | 0.206 | 0.186 | 0.203 | 0.188 | -0.020 | -0.015 | ud | ud | ud | ud | ud | ud | - | -0.15 |
| 18.03.2018 | 0.005 | -0.015 | -0.008 | -0.030 | -0.020 | -0.022 | ud | 46.5 | 90.0 | 23.5 | 151.4 | 76.5 | - | -0.02 |
| 19.03.2018 | 0.059 | 0.042 | 0.075 | 0.048 | -0.017 | -0.027 | ud | ud | ud | ud | ud | ud | -0.12 | -0.09 |
| 21.03.2018 | 0.051 | 0.038 | 0.078 | 0.039 | -0.014 | -0.039 | ud | ud | ud | ud | ud | ud | -0.06 | -0.07 |

While significant differences in growth rates at those stations were observed, *Synechococcus* populations declined at only three stations under FS and two stations in the -UV-B treatment, with mean half-life times (t_1/2_) of 35.4 and 32.9 h for FS and -UV-B, respectively (LRD_50_: 102.3 kJ) (Table 2). For pico-eukaryotes, cell concentrations decreased at six stations, regardless of the treatment, with mean t_1/2_ times of 12.6 and 38.8 h for the FS and -UV-B treatment, respectively (LRD_50_: 37.9 kJ).

We identified that natural *Synechococcus* populations near the surface (0.5 m) could be declining between 0 and 36% per day in summer UV-B conditions (Table 3). Population declines at the surface for January were calculated to be 0–21%, 3%, and 41%, respectively. In 8 m depth, *Synechococcus* population decay rates were <11% in summer and <4% in winter. Pico-eukaryotes displayed a wide variation in daily decay rates in surface water, ranging from 0–96% in summer and 0–82% in winter. In 8 m depth, cell abundance of pico-eukaryotes declined up to 32% per day in summer and 12% in winter.

**Table 3.** Theoretical daily population decays (in % of initial cell abundance) during July and January in various depths. Calculations are based on daily UV-B exposures (in kJ) reported by Overmans and Agusti (2020).

| Month | Depth (m) | UV-B (kJ d<sup>-1</sup>) | Cell decay (% d<sup>-1</sup>) |  |
| --- | --- | --- | --- | --- |
|  |  |  | *Synechococcus* sp. | Pico-eukaryotes |
| January | 0.5 | 19.2 | 0–21 | 0–82 |
|  | 2.0 | 11.4 | 0–13 | 0–63 |
|  | 8.0 | 1.4 | 0–2 | 0–12 |
| July | 0.5 | 36.6 | 0–36 | 0–96 |
|  | 2.0 | 23.7 | 0–25 | 0–88 |
|  | 8.0 | 4.3 | 0–5 | 0–32 |

### 4.2 Effects of UV-B on maximum photochemical efficiency (*F*_v_/*F*_m_)

The initial maximum photochemical efficiency (*F*_v_/*F*_m_) of phytoplankton communities at the start of the cruise experiments (T0) varied widely among experiments. Mean *F*_v_/*F*_m_ values ranged from 0.10 to 0.47 at the north (summer) and central (autumn) stations, respectively (Fig. 4). During summer, initial *F*_v_/*F*_m_ values were generally lowest in the morning, although some experiments showed exceptions to this pattern. In both light treatments, the phytoplankton community *F*_v_/*F*_m_ gradually decreased from T0 to T1 during summer and then recovered again by T2, reaching efficiencies similar to those measured in the morning. Notably, the community *F*_v_/*F*_m_ in the -UV-B treatment was higher at T1 than at T2 in the far north experiment, which was the only exception to this pattern.

**Figure 4.**
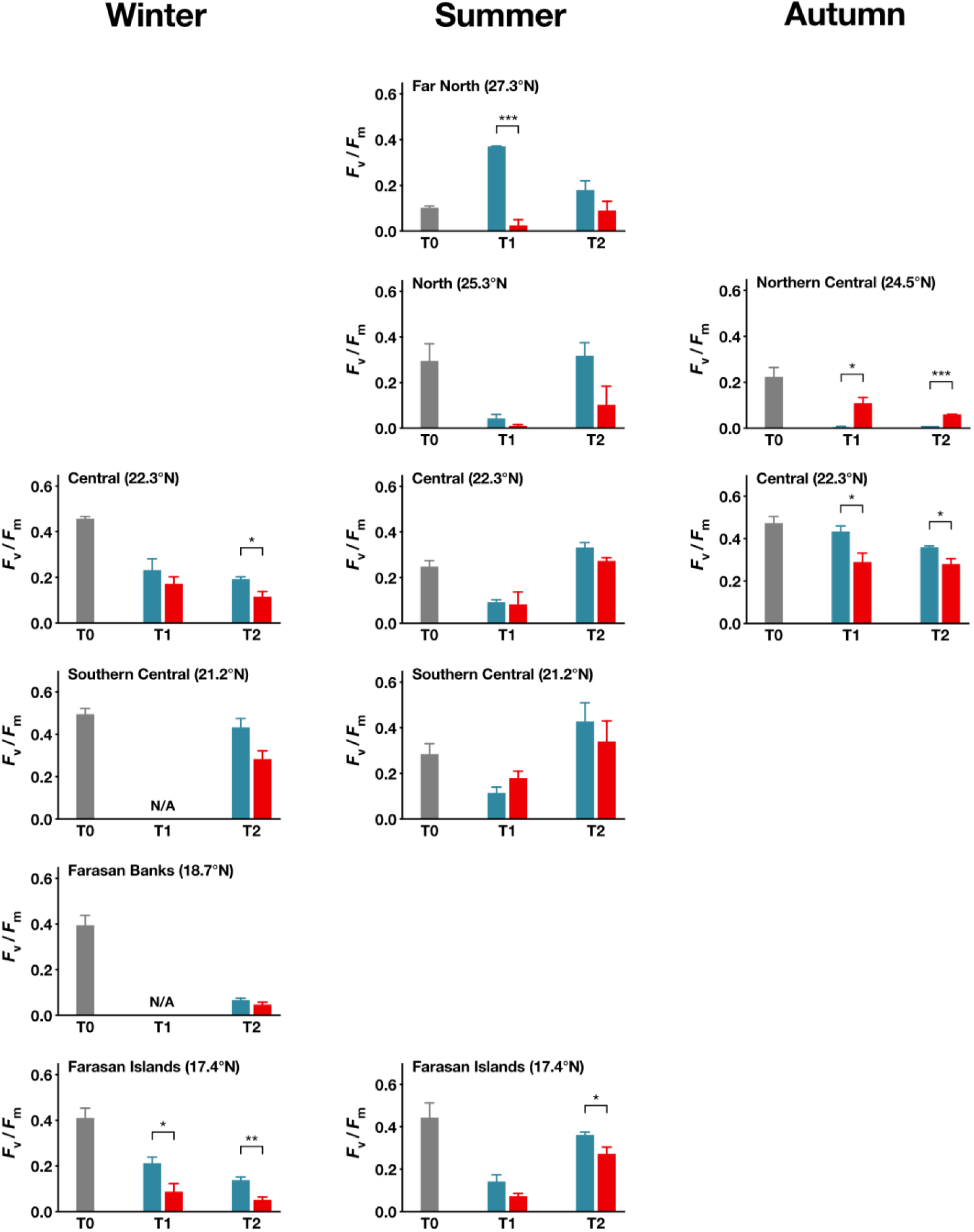
Maximum chlorophyll photochemical efficiency of photosystem II (*F*_v_/*F*_m_) of the phytoplankton community during the research cruise experiments in winter (left column), summer (middle column) and autumn (right column) from the northernmost station to the Farasan Islands in the far south of the basin (bottom row). For a description of the color-coding, see Fig. 2. Asterisks indicate significant differences in *F*_v_/*F*_m_ between treatments at a level of either *p* < 0.05 (*), < 0.01 (**), or < 0.001 (***).

Mean *F*_v_/*F*_m_ values were consistently higher in the UV-B reduction treatment than in the FS treatment at midday (T1) and in the late afternoon (T2) (mean ± SD Δ*F*_v_/*F*_m_: -0.08 ± 0.13 (T1), -0.08 ± 0.07 (T2); Table 2). When pooling measurements across all experiments, the mean *F*_v_/*F*_m_ was significantly higher in the -UV-B treatment (M= 0.18, SD= 0.14) than in the full sunlight (M= 0.12, SD= 0.09) at midday (T1) (t(60)= –2.35, *p* = 0.022) and early evening (T2) (M= 0.26, SD= 0.15 vs. M= 0.17, SD= 0.11, respectively; t(77)= –2.93, *p* = 0.0045). Statistically significant differences in mean *F*_v_/*F*_m_ between treatments were found only for specific experiments: winter experiments in the central Red Sea (T2) and the Farasan Islands (T1&2), summer experiments in the far north (T1) and the Farasan Islands (T2), and autumn experiment in the central sampling station (T1&2) (Fig. 4). At both sampling points during the experiment in the northern central in autumn, *F*_v_/*F*_m_ was significantly higher in the FS treatment than in the -UV-B treatment.

### 4.3 Role of environmental parameters in modulating UV-B sensitivity

No significant correlation between the growth rates (Δµ) and maximum photochemical efficiency (Δ*F*_v_/*F*_m_) of phytoplankton under UV-B-reduction conditions and their latitudinal origin were identified (Fig. 5), indicating that other environmental parameters may have a greater influence on these variables. However, a significant negative correlation was found between the Δµ of *Synechococcus* and SST (r = -0.61, *p* < 0.05), indicating that growth rates increased more under UV-B-reduction for cells from colder waters compared to those from warmer waters. The daily UV-B exposure was positively correlated with the Δµ of *Synechococcus*, suggesting population growth was more enhanced under UV-B-reduction in regions where daily UV-B doses were extremely high. No correlation was found between the Δµ of pico-eukaryotes and SST, UV-B exposure, or any other environmental parameter. Additionally, no significant correlations were observed between the Δ*F*_v_/*F*_m_ of the phytoplankton community and any environmental parameters.

**Figure 5.**
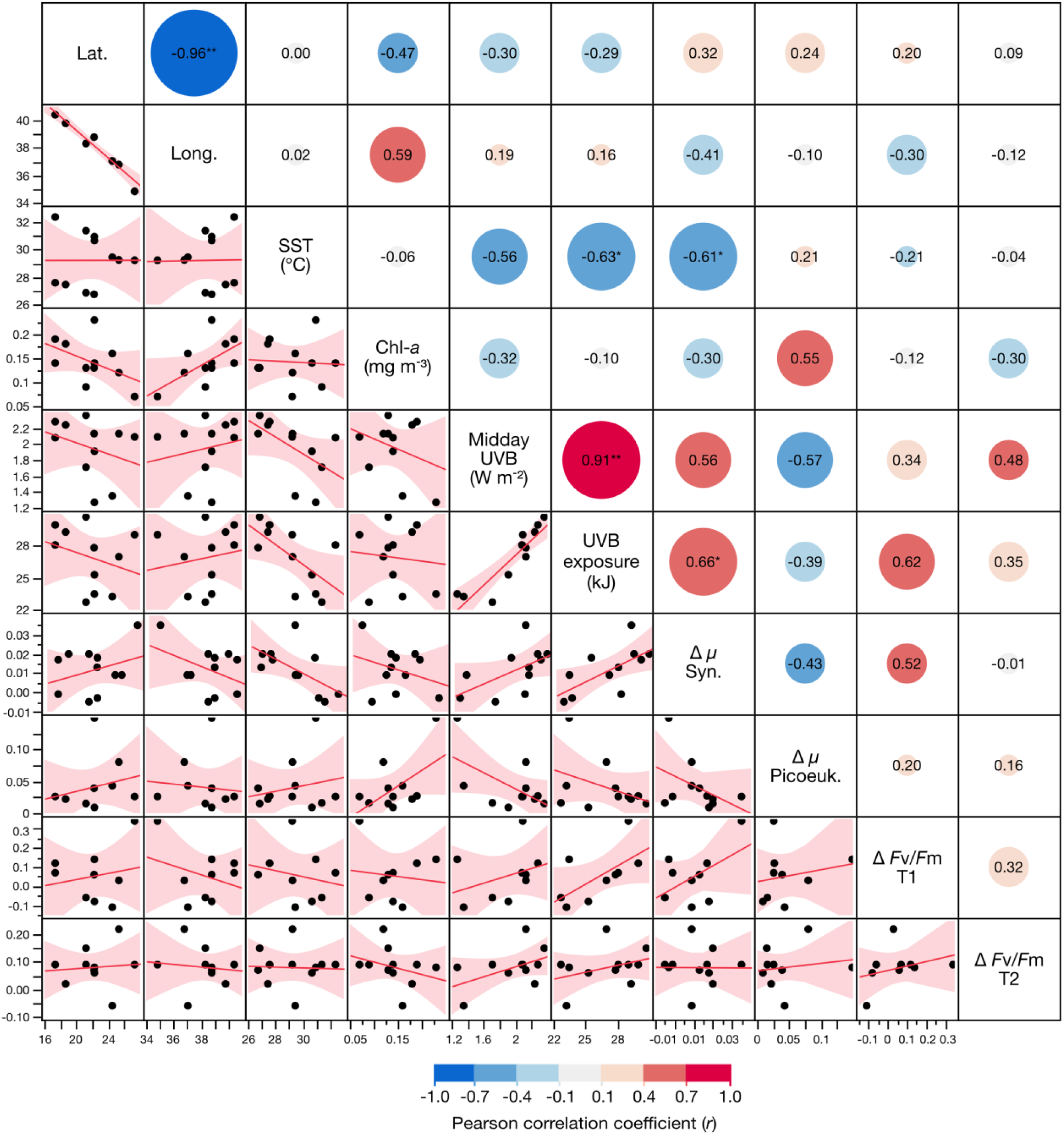
Results of the correlation analysis between various environmental parameters and changes in growth rates (Δµ) and the maximum photochemical efficiency (Δ*F*_v_/*F*_m_) of phytoplankton under UVB-reduction conditions. The red lines and shaded areas represent the correlation line and 95% CIs, respectively. The size of the circles and the values therein indicate the magnitude of the Pearson correlation coefficient (*r*). Asterisks indicate if the pairwise correlation analysis found *p* < 0.05(*) or *p* < 0.01(**).

## 5. Discussion

Overall, we identified that natural *Synechococcus* populations were highly resistant to UV-B-induced decay (from LRD_50_ 57.1 kJ to undetectable effects in 8 of 11 experiments), unlike pico-eukaryotes (from LRD_50_: 7.8 kJ to undetectable effects in 5 of 11 experiments). Across experiments, the reduction in phytoplankton community *F*_v_/*F*_m_ due to UV-B exposure (Δ*F*_v_/*F*_m_) was small (-0.08 ± 0.13 around noon and -0.08 ± 0.07 in the early evening) but significant.

We could observe that natural Red Sea surface water communities, dominated by *Synechococcus*, exhibited relatively minor UV-B-induced reduction in their maximum photosynthetic efficiency (Δ*F*_v_/*F*_m_) of -0.08 ± 0.13 (Mean ± SD) around noon. This result highlights that the continued exposure of natural communities to high UV-B intensities has likely led to selection of well-acclimatized and UV-resistant assemblages in surface waters, as previously suggested for Antarctic phytoplankton (Boyd et al., 2000; Seckbach and Oren, 2007).

The smallest cells, such as the pico-cyanobacteria *Synechococcus*, exhibited the highest resistance to UV-B-induced damage despite their high surface-to-volume ratio. However, as cell size decreases, the cost/benefit ratio of non-optical defence strategies such as antioxidants, repair, and reactivation processes significantly improve (Raven, 1991). Therefore, the observed high resistance of *Synechococcus* to UV-B might be because pico-cyanobacteria possess several non-optical defence mechanisms against UV-B radiation, including multiple copies of their genome, adaptive mutagenesis, or the synthesis of ROS-quenching enzymes and non-enzymatic antioxidant compounds (Kumar et al., 2016; Prabha and Kulandaivelu, 2002). Marine *Synechococcus* strains can also integrate a more resistant D2 protein into PSII when exposed to UV (Mella-Flores et al., 2012), resulting in a substantial decrease in sensitivity to inhibition of quantum yield of PSII and photosynthesis by subsequent UV exposure (Fragoso et al., 2014; Neale et al., 2014). Additionally, the activity of plastid alternative oxidase (PTOX) has also been shown to be important in limiting damage to PSII (Grossman et al., 2010). Small cells such as pico-cyanobacteria are also characterized by fast metabolic rates and acclimation kinetics (Peters, 2012; Villafañe et al., 2004), thereby enabling a rapid repair of UV-damaged cellular structures such as DNA or PSII (Key et al., 2010; Mella-Flores et al., 2012; Six et al., 2007). While some studies found that UV-B sensitivity is strongly dependent on cell size (Li and Gao, 2013; Wu et al., 2015), other cellular characteristics such as cell morphology, the distribution of organelles, DNA structure, and antioxidants and repair capabilities may be of greater importance (He and Hader, 2002; Helbling et al., 1992; Wängberg et al., 1999). The diversity of defence strategies may explain the low vulnerability to UV-B of natural *Synechococcus* populations reported here and account for the previously described high resistance to UV-B of *Synechococcus* strains in culture (Mella-Flores et al., 2012), as well as of natural assemblages from the Mediterranean Sea (Sommaruga et al., 2005) and the tropical Atlantic Ocean (Boelen et al., 2000).

However, our results that *Synechococcus* populations were still growing under full sunlight at most stations (µ: 0.044 ± 0.019 h^−1^, mean ± SE) are contrary to previous findings suggesting a moderate to high decay of natural *Synechococcus* populations due to UV-B exposure, where *Synechococcus* cell decay of in tropical Atlantic waters was dramatically increased when the cells were exposed to solar PAR + UVR (µ: -0.021 ± 0.008 h^−1^, mean ± SE) as opposed to PAR-only (Llabres and Agusti, 2006). Similar results were obtained for natural communities from the Mediterranean Sea, where *Synechococcus* exhibited a high sensitivity to UV-B, as determined by considerably shorter half-life times under full sunlight (t_1/2_: 9.9 h) than in the absence of UV-B (t_1/2_: 29.8 h), and an average lethal UV-B dose (LRD_50_) of 30.4 kJ, which is considerably lower than in the present study (LRD_50_: 57.1 kJ to undetectable) (Llabres et al., 2010). However, we could only determine LRD_50_ values in three experiments since populations did not decline during the other eight experiments, which suggests that Red Sea *Synechococcus* populations are considerably more resistant to UV-B than those in the Mediterranean. The much lower UV-B sensitivity reported here could stem from the difference in the two experimental setups, since Llabres et al. (2010) additionally shielded the cells from UV-A wavelengths, which are known to aid repair processes (Häder et al., 2015). Additionally, it is noteworthy that Llabres et al. (2010) specifically determined the viability of the cells, whereas we calculated the decay rates and LRD_50_ based on cell abundance, which might, at least in part, account for the large discrepancy between the two results.

Moreover, the observed discrepancy between the findings of Llabres et al. (2010) and our results might be due to genetic differences between the populations that result in varying sensitivities, or more specifically, the *Synechococcus* populations native to the Mediterranean Sea may belong to a different clade than the ones in Red Sea waters. For example, it has been identified that *Synechococcus* in the Mediterranean Sea primarily belongs to clade III, which is more commonly found at mid-latitudes (Farrant et al., 2016; Lee et al., 2019; Mella-Flores et al., 2011). In contrast, the predominant clade of *Synechococcus* in the Red Sea is clade II (Fuller et al., 2005; Lee et al., 2019), and clade II-a in particular (Farrant et al., 2016), which is known to be the prevailing clade in the warm and intensely illuminated waters of the tropics (Ahlgren and Rocap, 2012; Fuller et al., 2006; Pittera et al., 2014). While *Synechococcus* clade II is the dominant clade throughout the water column, other clades (e.g., clades III, VII, VIII) can be found in less well-lit layers of the photic zone (Fuller et al., 2003; Fuller et al., 2005; Lindell and Post, 1995). Our *Synechococcus* strain from the Gulf of Aqaba (2383) belongs to the uncommon clade VIII, while the central Red Sea strain (CB1) is a mix-population predominantly composed of cells from the common clade IIa. Since the water in the present study was collected close to the surface (5 m depth), it is apparent why we predominantly sampled *Synechococcus* clade II, which is well-adapted to the intense UV-B conditions prevalent in shallow depths. This sampling strategy would also explain why we did not detect the cyanobacterium *Prochlorococcus* in any of our samples, which is known to be vulnerable to intense irradiation and ultraviolet radiation in particular (Llabres and Agusti, 2006; Llabres et al., 2010; Mella-Flores et al., 2012), and not commonly found in Red Sea surface waters but instead abundant in deep or intermediate depths of the photic zone (Coello-Camba and Agusti, 2021; Dishon et al., 2012a).

Our results indicate that the observed variabilities in the community photoinhibition and decay rates of *Synechococcus* and pico-eukaryotes between experiments could be the result of distinct light histories of the cells at the various stations. Previous studies have suggested that historic light conditions can considerably affect the vulnerability to UV-B, with cells that had previously been exposed to UV-B exhibiting a higher resistance upon re-exposure, as shown for pico-cyanobacteria, diatoms, and dinoflagellates (Guan and Gao, 2008; Häder et al., 2014; Llabres et al., 2010). Therefore, the observed differences in UV-B sensitivity could also be attributed to genotypic differences and adaptations of natural phytoplankton communities to the specific environmental conditions prevailing at each station. Since the Red Sea is characterized by distinct gradients in its UV penetration properties (Overmans and Agusti, 2019) and vertical mixing (i.e., mixed layer depths) (Abdulla et al., 2018), the spatial variation in the UV-B sensitivity of phytoplankton may be at least in part modulated by the light conditions. While we did not find a significant relationship between Δµ and latitude for either *Synechococcus* or pico-eukaryotes (see Fig. 5), the vastly different light histories of the cultured and natural *Synechococcus* could explain why the cultured strains showed a higher UV-B sensitivity. A further factor that influences the vulnerability of Red Sea phytoplankton to UV-B is the availability of nutrients since a nutrient limitation can increase the sensitivity of phytoplankton to UV-B (Litchman et al., 2002; Neale and Kieber, 2000; Xenopoulos et al., 2002). While there was no significant correlation between Chl *a* concentration, representative of trophic degree, and Δµ of *Synechococcus* and pico-eukaryotes, a study from the North Atlantic Ocean reported that the UV sensitivity of pico-eukaryotes was closely coupled with Chl *a* concentration (Agusti and Llabres, 2007). In the latter study, however, experiments were performed in several locations of varying trophic states (Chl *a*: 0.09–4.26 mg m^−3^), while our experiments were exclusively performed in highly oligotrophic waters (Chl *a*: 0.14–0.23 mg m^−3^), which may explain the contrasting results.

We did, however, find a significant negative correlation between SST and the change of *Synechococcus* growth rates under UV-B reduction (r = -0.61, *p* < 0.05) (Fig. 5), which implies that changes in growth were higher in cooler waters less pronounced in the warmest waters under UV-B reduction. We also found a weak negative correlation between temperature and Δ*F*_v_/*F*_m_ of the phytoplankton community at T1 (r = -0.21). Both findings are in agreement with several reports suggesting that elevated temperatures can accelerate growth and cellular repair mechanisms, and therefore increase UV-B resistance (Bouchard et al., 2006; Halac et al., 2014; Helbling et al., 2011). However, this relation only applies within the thermal limits of an organism, and if temperatures are beyond the thermal optimum, the detrimental effects of excessive temperature can interact synergistically with harmful UV-B, resulting in multiplicative impacts in a wide range of organisms (Jin et al., 2019). Here, the UV-B sensitivity of neither *Synechococcus* nor pico-eukaryotes exhibited a latitudinal gradient (see Fig. 5), possibly because the cells are not experiencing as much UV-B-induced stress since the northern Red Sea receives lower incident UV-B irradiances than southern regions (Dishon et al., 2012b; Overmans and Agusti, 2020).

Moreover, several reports have highlighted that the effect of UV-B is strongly dependent on the spectral balance (Quesada and Vincent, 1997). It has been shown that UV-B impacts on autotrophs are exacerbated under unnaturally high ratios of UV-B: UV-A and UV-B: PAR (Krizek, 2004; Ryan et al., 2012), which implies that any experimental results that were obtained using artificial light sources with unnatural spectra cannot necessarily be extrapolated to assess responses of natural communities reliably. It should also be considered that any damage caused by UV-B during the incubation may have been slightly exaggerated due to the reduction of UV-A radiation, especially since UV-A was found to have several beneficial effects on marine organisms when received in moderate exposures (Häder et al., 2015). For example, UV-A is involved in the photoreactivation of DNA and the activation of repair processes (Li et al., 2011; Quesada et al., 1995; Sinha and Häder, 2014), which highlights the significance of spectral composition, and more specifically the presence of UV-A, when evaluating UV-B impacts on phytoplankton.

## 6. Conclusions

Our study reveals that Red Sea *Synechococcus* sp. populations are highly resistant to UV-B exposure, while natural pico-eukaryotes show higher susceptibility, particularly during the summer. We identified that the decay rates of *Synechococcus* and pico-eukaryotes under full sunlight did not correlate with latitude or incubation temperature. However, when UV-B was reduced, *Synechococcus* from colder waters showed considerably increased growth rates. We also observed considerable intra-specific variability in UV-B resistance, which was likely driven by differences in the light history, spectral composition, and nutrient availability. In summary, our study indicates that many of the Red Sea phytoplankton, particularly *Synechococcus*, exhibit higher UV-B resistance compared to phytoplankton from other basins, highlighting their adaptation to the extreme UV-B exposure in these waters.

## Author contributions

SA and SO conceptualized the study and analysed the data. SO performed the experiments and collected the data. SO generated tables and figures. SO prepared the first draft of the manuscript, but both authors contributed substantially to subsequent versions, including the final draft.

## Acknowledgements

We would like to express special thanks to Juan Martinez Ayala, Veronica Chaidez and Daffne Celeste López Sandoval from the Biological Oceanography Laboratory, and the staff of the Coastal and Marine Resources (CMR) Core Lab at King Abdullah University of Science and Technology (KAUST), and the crews of the R/V Thuwal and R/V Al Azizi for their support and assistance during the fieldwork and laboratory analyses. The research reported in this publication was supported by the KAUST baseline funding of S. Agustí under award number BAS/1/1072-01-01 and the KAUST RSRC Center Competitive Fund (CCF).

## Notes

### Competing Interest Statement

The authors have declared no competing interest.

