## Supplementary material for "Red Sea *Synechococcus* exhibits greater resilience to UV-B radiation than pico-eukaryotic phytoplankton": Suppl. Fig. 1

**<u>Contents of this file</u>**

Supplementary Figure 1

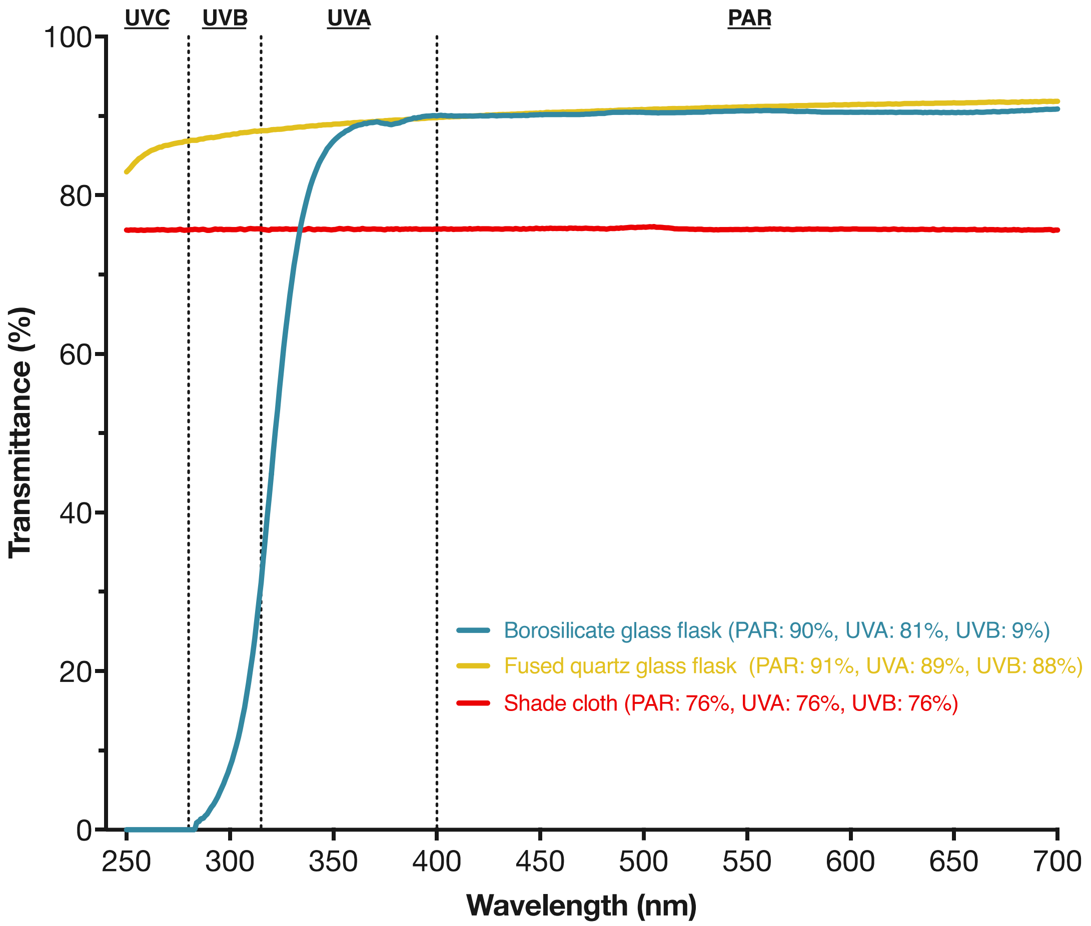


**Supplementary Figure 1.** Transmission spectra of different materials that were used as part of the phytoplankton experiments. Spectra were measured using a UV/VIS/NIR spectrophotometer (Model Lambda 1050; PerkinElmer, USA).
